# Self-supervised deep learning of gene-gene interactions for improved gene expression recovery

**DOI:** 10.1101/2023.03.10.532124

**Authors:** Qingyue Wei, Md Tauhidul Islam, Yuyin Zhou, Lei Xing

## Abstract

Single-cell RNA sequencing (scRNA-seq) has emerged as a powerful tool to gain biological insights at the cellular level. However, due to technical limitations of the existing sequencing technologies, low gene expression values are often omitted, leading to inaccurate gene counts. The available methods, including state-of-the-art deep learning techniques, are incapable of imputing the gene expressions reliably because of the lack of a mechanism to explicitly consider the underlying biological knowledge of the system. Here we tackle the problem in two steps to exploit the gene-gene interactions of the system: (i) we reposition the genes in such a way that their spatial configuration reflects their interactive relationships; and (ii) we use a self-supervised 2D convolutional neural network to extract the contextual features of the interactions from the spatially configured genes and impute the omitted values. Extensive experiments with both simulated and experimental scRNA-seq datasets are carried out to demonstrate the superior performance of the proposed strategy against the existing imputation methods.

## Introduction

The success of transcriptomic studies, such as differential expression analysis^1–3^, cell subpopulation identification^4–6^, cell trajectory reconstruction^7–9^, and alternative splicing detection^10,11^ depends critically on the accuracy of the gene expression counts. In practice, gene expression data often suffer from low transcript capture efficiency and technical noise, leading to inaccurate gene counts. Thus, recovery of the expression values of the genes using computational techniques is critical for the downstream applications.

Broadly, gene expression recovery techniques can be divided into three main categories: 1) probabilistic models, 2) nearest neighbor-based techniques, and 3) deep learning methods. SAVER^12^, SAVER-X^13^, BayNorm^14^, and scImpute^15^ are the prominent techniques of the first group. SAVER^12^ and BayNorm^14^ are two Bayesian approaches that use the gene-gene relationships to estimate the expression levels of the omitted genes. scImpute^15^ identifies and imputes the dropout gene expression values by applying a known statistical model. The second group uses the information from the neighboring genes to interpolate the omitted gene expressions. MAGIC^16^ explores information across similar cells and utilizes a Markov matrix to denoise the gene expression data while imputing the dropouts. DrImpute^17^ is another method of this group, which integrates information from similar cells and adopts expression averaging for data recovery. AutoImpute^18^, DeepImpute^19^, scVI^20^, and scScope^21^ are the representative methods of the third category. These methods use different network architectures such as autoencoder (AutoImpute^18^), multilayer perceptron (DeepImpute^19^), and recurrent neural network (scScope^21^), to extract information from similar cells and genes for data recovery. These existing methods suffer from either accuracy or computational efficiency or both, which limits their practical applications. The first group (*e.g*. SAVER) assumes a specific distribution of gene expression data which may not be sufficiently accurate in practical scenarios. The second type of methods such as MAGIC relies on expression averaging for imputation, which may result in removal of the variability in gene expression. Deep learning techniques have limited receptive field size and may lead to suboptimal results due to their difficulties in capturing the long-range relationship between cells and genes.

It is well known that the gene-gene interaction patterns are unique to biological systems and present discriminative signatures of the cells involved. Here we leverage the interactive information and establish a transform-and-conquer expression recovery (TCER) strategy to tackle the gene imputation problem. First, we transform the gene expression data into an image format, referred to as the GenoMap, based on the interactions among the genes. In this step, the gene-gene interactions of the system are configured by placing the genes in such a way that the genes interacting strongly are close to each other. This transformation maps the gene-gene interactions into configured format and enables a deep neural network (DNN) to exploit the interactions more effectively. Thus, the method mitigates the the problem of limited receptive field of conventional deep learning approaches and facilitates full exploitation of the gene-gene relationships.

To extract deep interaction information from a GenoMap for the recovery of missing expression values, a novel encoderdecoder architecture, referred to as expression recovery network (ER-Net), is designed. In ER-Net, we include three cascaded Deformable Fusion Attention (DFA) modules between an encoder and a decoder. Each DFA module includes a deformable convolution layer so that its kernel shape can be adaptively adjusted according to the input feature maps. The deformable kernels enable the network to explore gene-gene relationships more flexibly and enhance the capability for the network to discover the underlying patterns in GenoMaps. Additionally, the network uses a dual-attention (channel-wise and pixel-wise attention) mechanism to adaptively assign higher values to more important channels and positions for high-performance expression recovery.

Extensive experiments on the simulated scRNA-seq data and six real-world scRNA-seq datasets are performed. We demonstrate that the proposed method substantially outperforms the existing ones in terms of imputation accuracy. We show that the recovered data using the proposed technique also yields the best outcomes of cell clustering and trajectory analyses.

## Results

### TCER enables reliable gene imputations

We use the Splatter simulator^22^ to simulate a reference scRNA-Seq data based on a gamma-Poisson distribution for 10000 cells each with 2400 genes. We set the number of cell groups to 5 with a probability of 0.2 for each group. To imitate the experimental scRNA-Seq data acquisition process, we sample the reference data following ref.^12^ to generate an observation dataset with 1% efficiency loss. We compare the performance of TCER with two existing gene imputation methods, scVI^20^ and MAGIC^16^. In Fig. 4, we show t-SNE^23^ and UMAP^24^ visualization of the reference data (first column), observation data (second column), and results from TCER (third column), scVI (fourth column), and MAGIC (fifth column). It is obvious that our method shows better-separated clusters when compared to the results directly from the observation data and imputed data from scVI and MAGIC. scVI performs relatively better than MAGIC. We calculate the clustering accuracy and quality indices to quantitatively evaluate the clusters resulted from different methods. As shown in Fig. 4, TCER greatly outperforms other methods.

**Figure 1.**
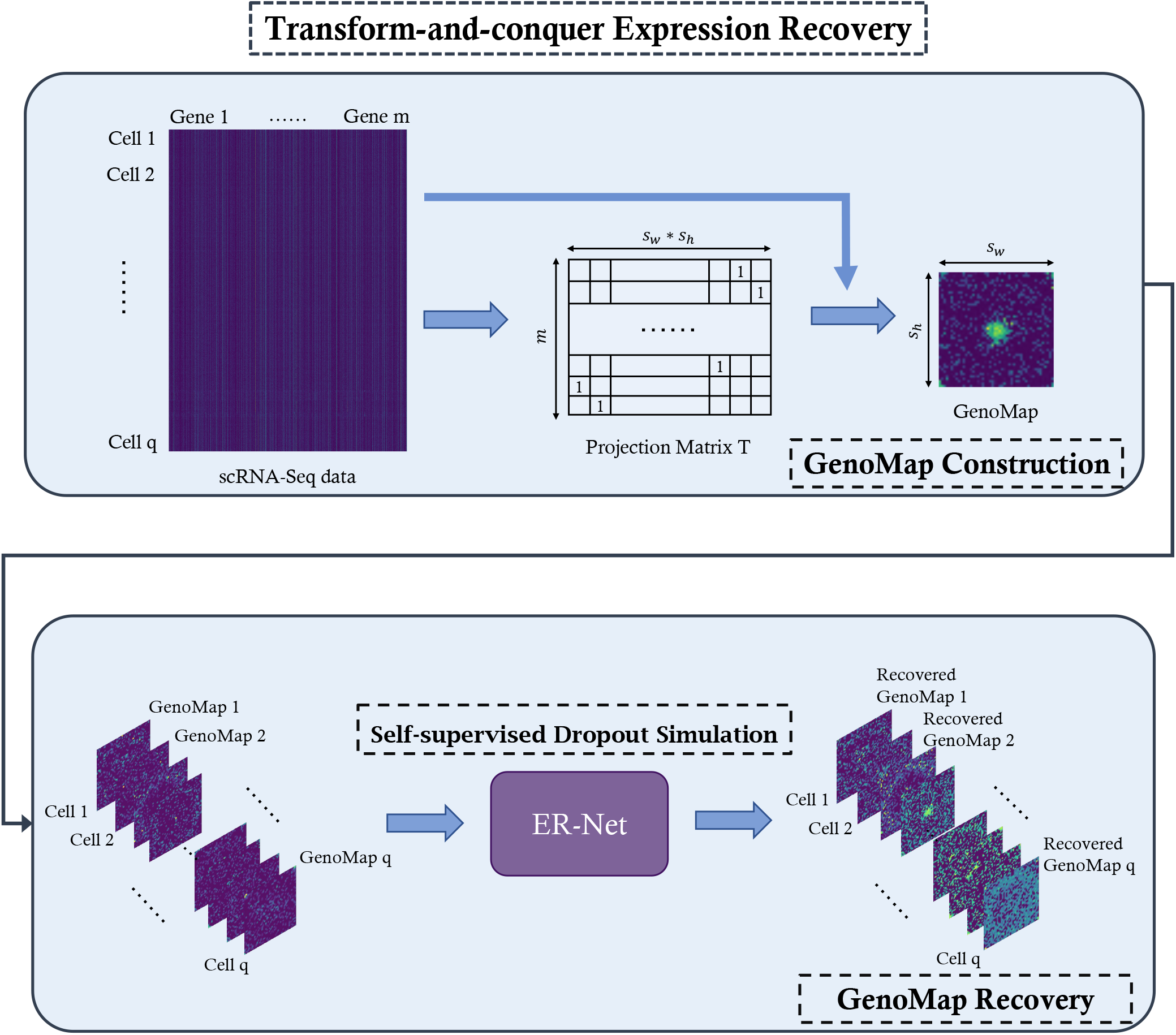
Pipeline of the proposed TCER. 1D gene expression data is first converted into an image format where the gene-gene interactions are reflected naturally in the spatial configuration of GenoMap. A dropout simulation strategy is then applied to simulate the dropout events where non-zero values are randomly masked. Last, the proposed ER-Net is employed to impute the masked GenoMap.

**Figure 2.**
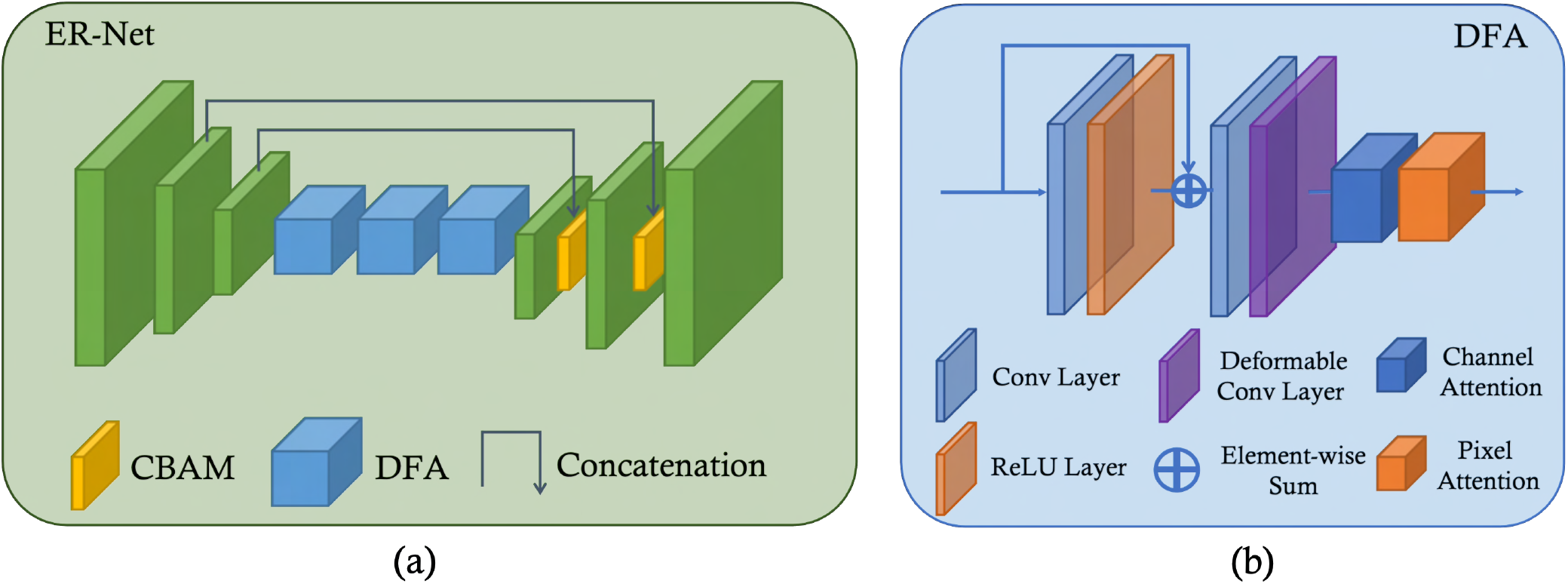
Network structure of ER-Net. (a) ER-Net follows an encoder-decoder structure and employs three cascaded DFA module with deformable convolution to extract both local and global features of gene-gene interactions. b) Detailed structure of the proposed Deformable Fusion Attention (DFA) module.

**Figure 3.**
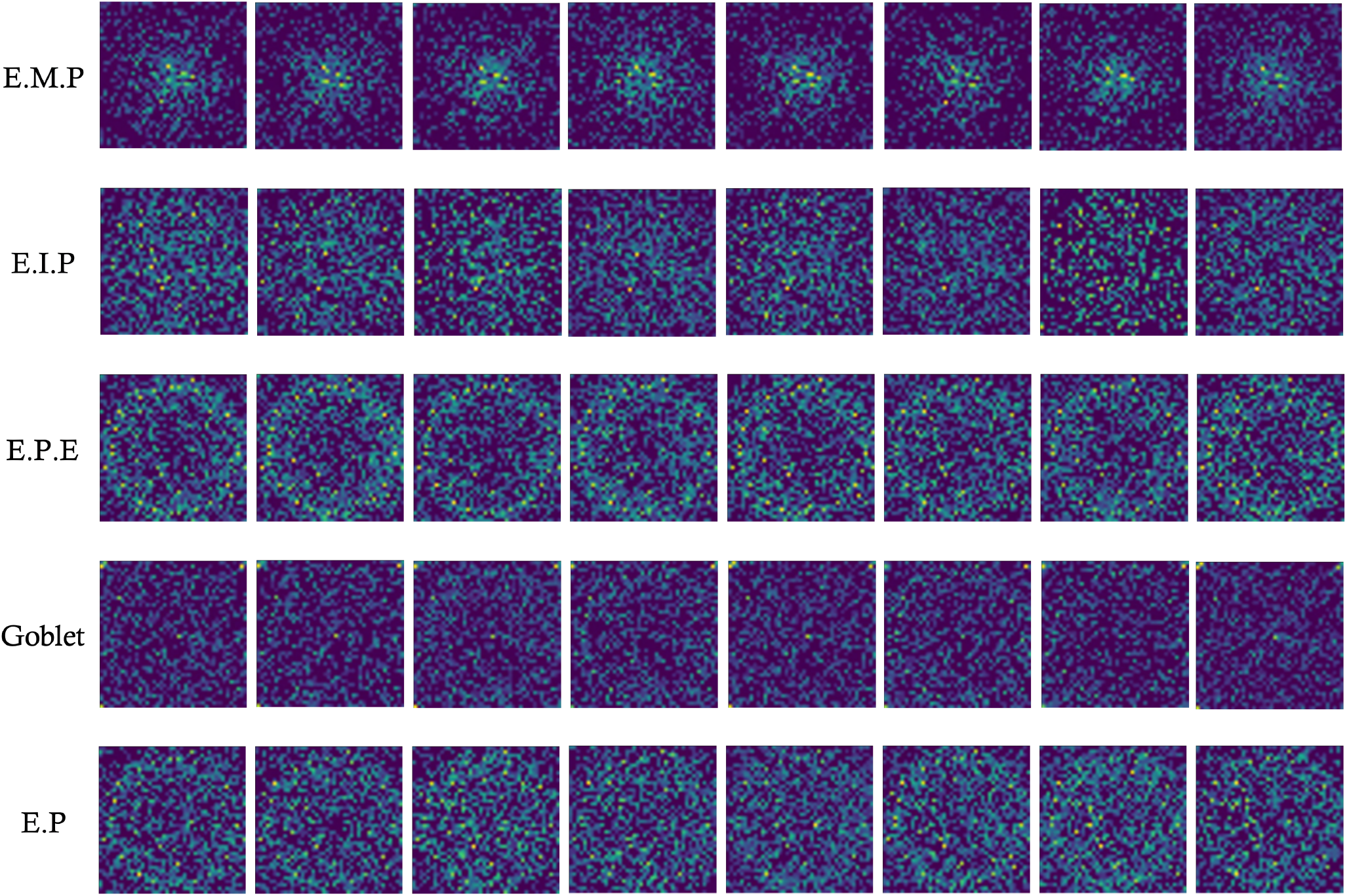
Visualization of GenoMaps of different cell types for mouse intestinal epithelium dataset. Each row represents GenoMap from 8 different cells in the same cell type.

**Figure 4.**
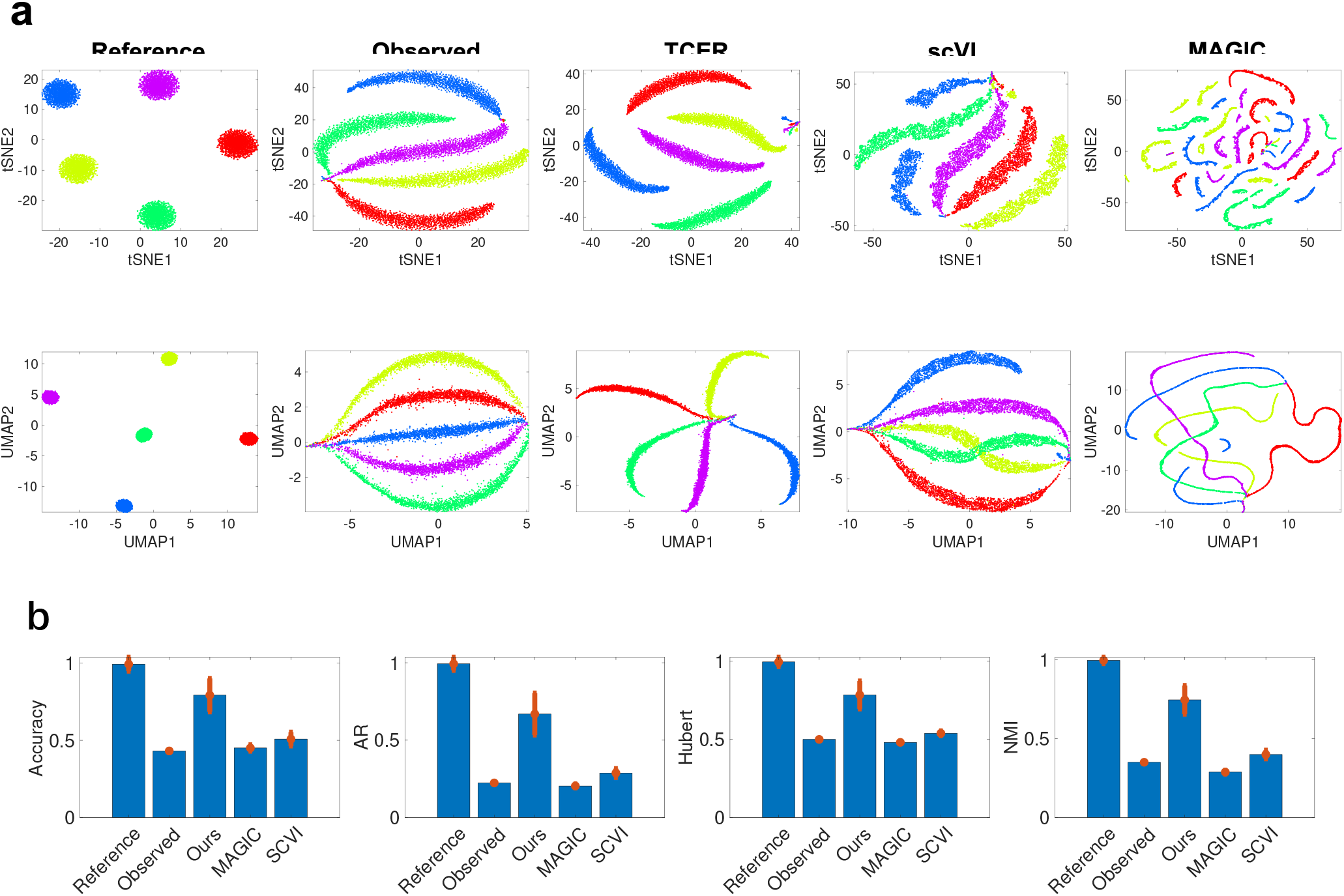
Analyses of simulation dataset. (a) tSNE and UMAP visualizations of reference (first column), observed data (second column), and imputed data by TCER (third column), scVI (fourth column) and MAGIC (fifth column). (b) The clustering accuracy and cluster quality indices for UMAP visualizations of the reference and observed data, and imputed data using different methods.

TCER also outperforms other methods in real-world settings. To demonstrate this, we conduct experiments using four experimental scRNA-seq datasets: 1) mouse intestinal epithelium^25^, 2) engineered 3D neural tissues^26^, 3) mammalian brains^27^, and 4) cellular taxonomy^28^. Details of the datasets are described in the Methods section. For each dataset, we first select cells and genes with high expression levels and use them as the reference data. We then sample the reference data using a negative binomial model^12^ to generate observation datasets at different efficiency losses. In Fig. 5, we show the GenoMaps of the reference and observation datasets, and the recovered GenoMaps by TCER. The recovered GenoMaps correlate strongly to the reference ones.

**Figure 5.**
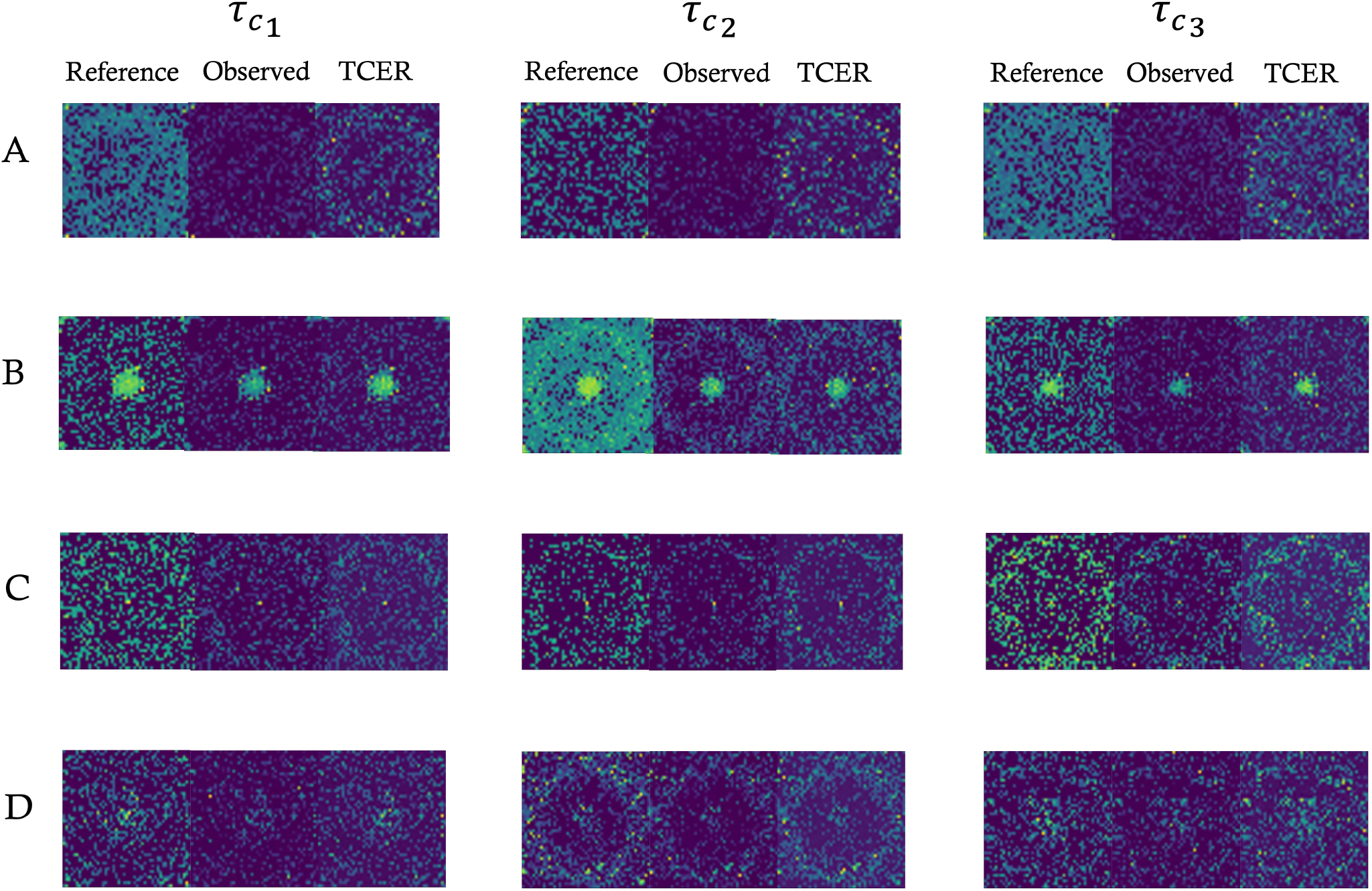
GenoMaps of the reference, observed and recovered data from different datasets sampled at different efficienciesEach row displays samples from the same dataset with different efficiency loss. Datasets include (A) mouse intestinal epithelium, (B) 3D neural tissue data, (C) mammalian brain, (D) cellular taxonomy. And *τ*_*c* 1_ represents 0.75% efficiency, *τ*_*c*2_ indicates 0.5% efficiency while *τ*_*c*3_ means 0.4% efficiency.

We present the UMAP visualizations of original data and data imputed by different methods for all four datasets in Fig. 6 It is seen that, compared to the results from other methods, UMAP visualizations of our imputation results correlate better to the reference data (the first column) with superior clustering quality. We note that data distortions are observed in all four datasets.

**Figure 6.**
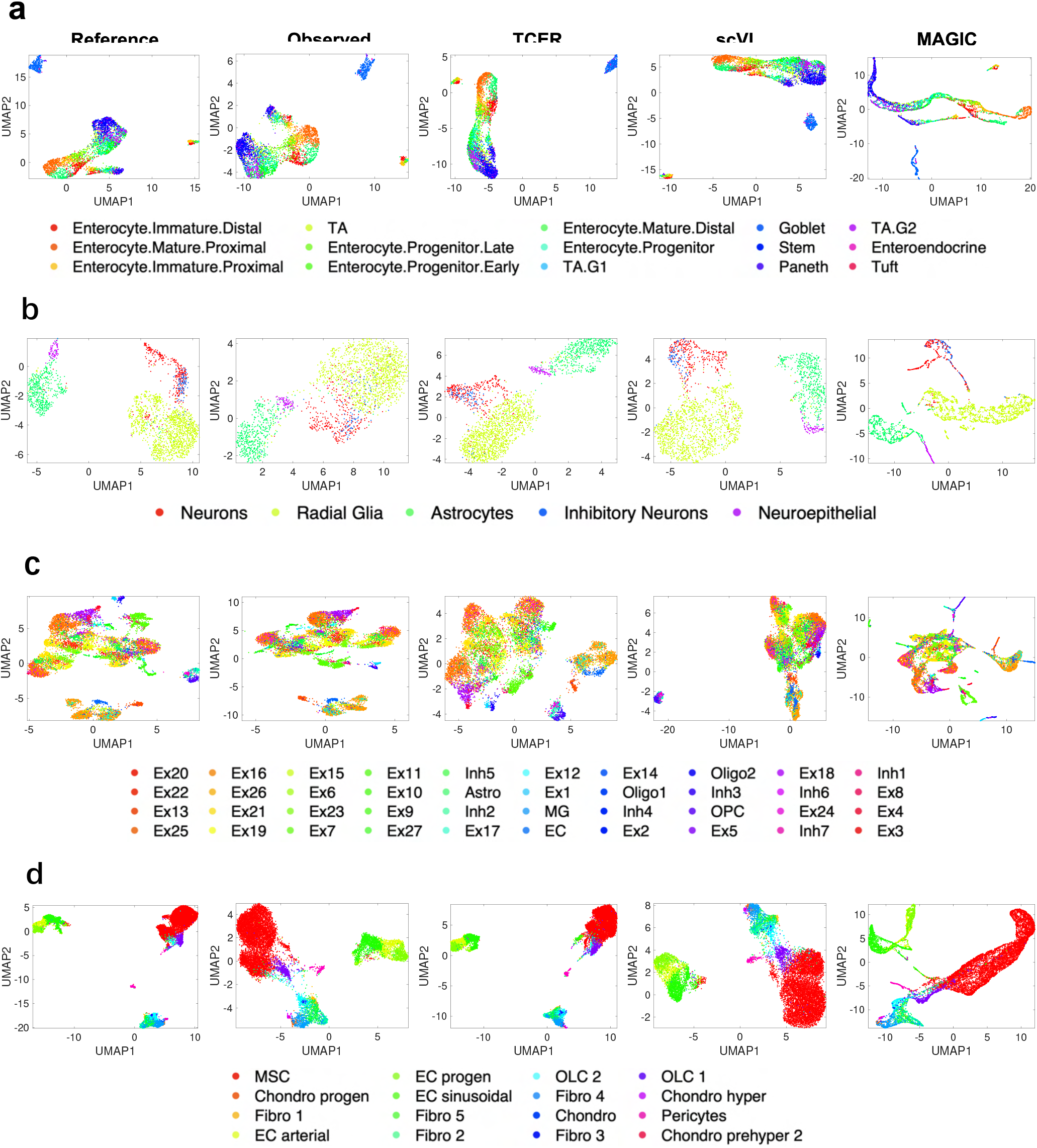
UMAP visualizations of the reference (first column) and observed data (second column), and the imputed results from TGER, scVI, and MAGIC. Here, (a) mouse intestinal epithelium is sampled at 0.75% efficiency, (b) 3D neural tissue data is sampled at 0.5% efficiency, (c) mammalian brain is sampled at 0.4% efficiency, and (d) cellular taxonomy data is sampled at 0.4% efficiency.

The Pearson correlation results from different methods are shown in Fig. 7 It is seen that TCER yields the best performance for all datasets with more than 6% improvement in Pearson coefficients as compared to that obtained from the observed datasets directly. It is worth noting that the other three methods lead to Pearson coefficients lower than that from the original data without any imputation. For example, for an efficiency loss of 0.4% for mammalian brains dataset, Pearson coefficients resulted from original data, TCER-, MAGIC- and scVI-imputed data are 0.7615, and 0.8005, 0.3326, and 0.2469, respectively.

**Figure 7.**
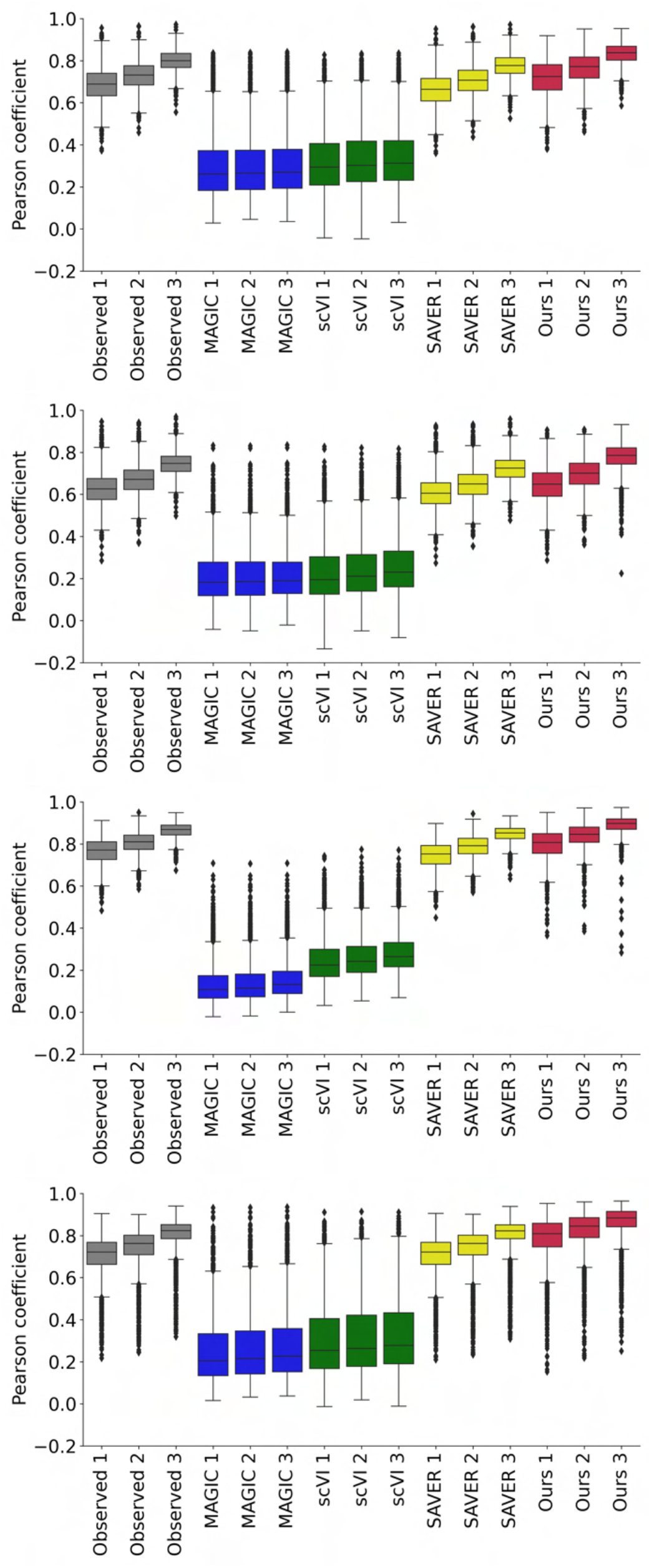
Box plots of Pearson coefficient between the reference and imputed data from different methods for mouse intestinal epithelium (first row), 3D neural tissue data (second row), mammalian brain (third row), cellular taxonomy (fourth row). 1-3 shown in x-axis indicate the sampling efficiency 0.4%, 0.5%, and 0.75%.

### TCER improves the accuracy of cell trajectory analysis

Cell trajectory analysis is widely used to understand the cellular differentiation process and plays an important role in the study of embryo and organ developments. Here, we use two experimental datasets to demonstrate that TCER can restore the missing expressions of trajectory data and facilitates the analysis of cell trajectories. The datasets are from 1) human Embryonic Stem (ES) cells differentiated to embryoid bodies (EBs)^29^ and 2) early zebrafish development^30^. Details of each dataset can be found in the Methods section. We employ the well established PHATE^31^ to embed the high-dimensional data for trajectory visualization and analysis. The PHATE results for imputed data from different approaches are shown in Fig. 8. It is seen that our imputed results (third column) contain continuous cell trajectory patterns that closely resemble the reference data (first column), whereas the observation data (second column) and imputed results from scVI (fourth column) and MAGIC (fifth column) show distortions in the trajectories. In Fig. 9, we show the Pearson coefficients with respect to the reference dataset for all the cases. Again, TCER method achieves the best Pearson coefficients for both datasets.

**Figure 8.**
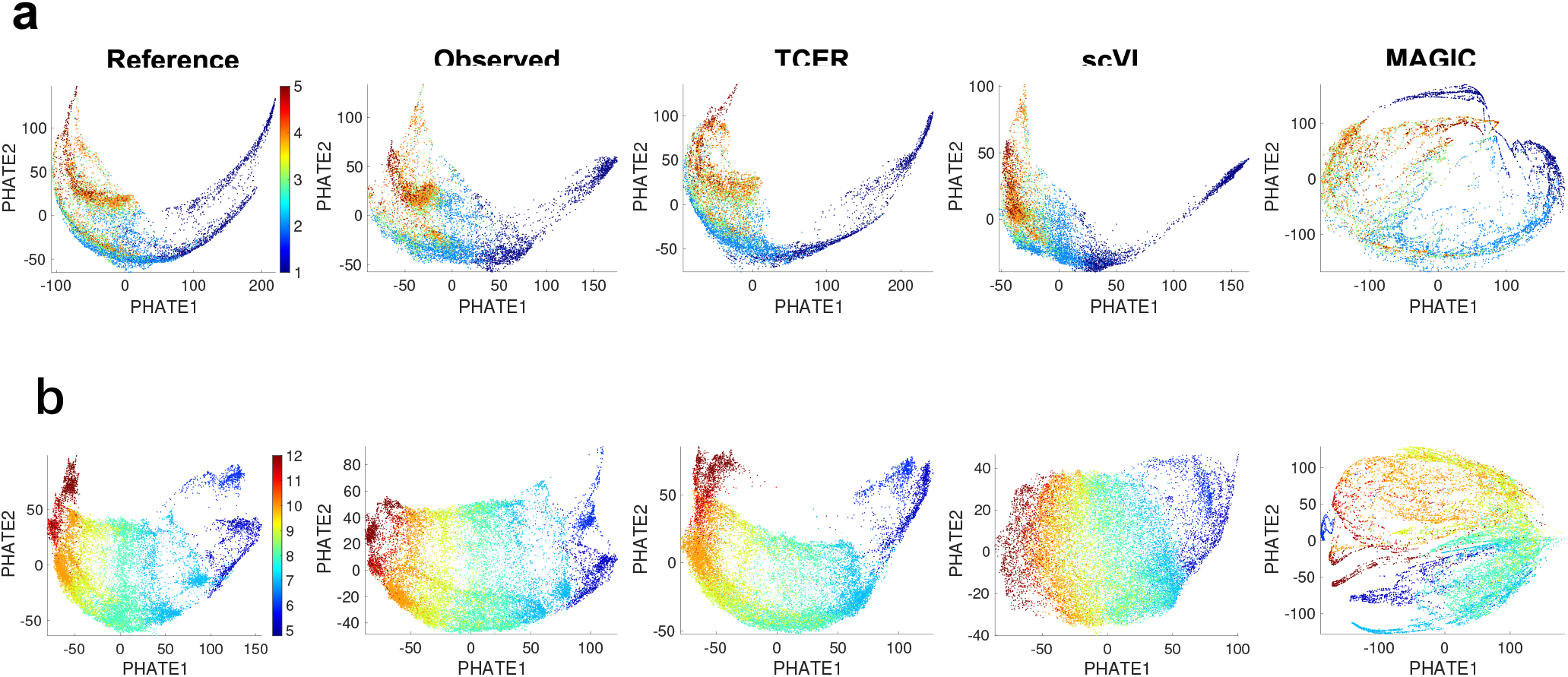
PHATE visualizations of reference and observed data, and imputed results from TCER, scVI, and MAGIC for (a) EB differentiation data sampled at 0.75% efficiency and (b) zebrafish development data sampled at 0.5% efficiency. The colorbar for (a) indicates 1-(0-3 days), 2-(6-9 days), 3-(12-15 days), 4-(18-21 days) and 5-(24-27 days) and (b) shows the hpf (hours post fertilization).

**Figure 9.**
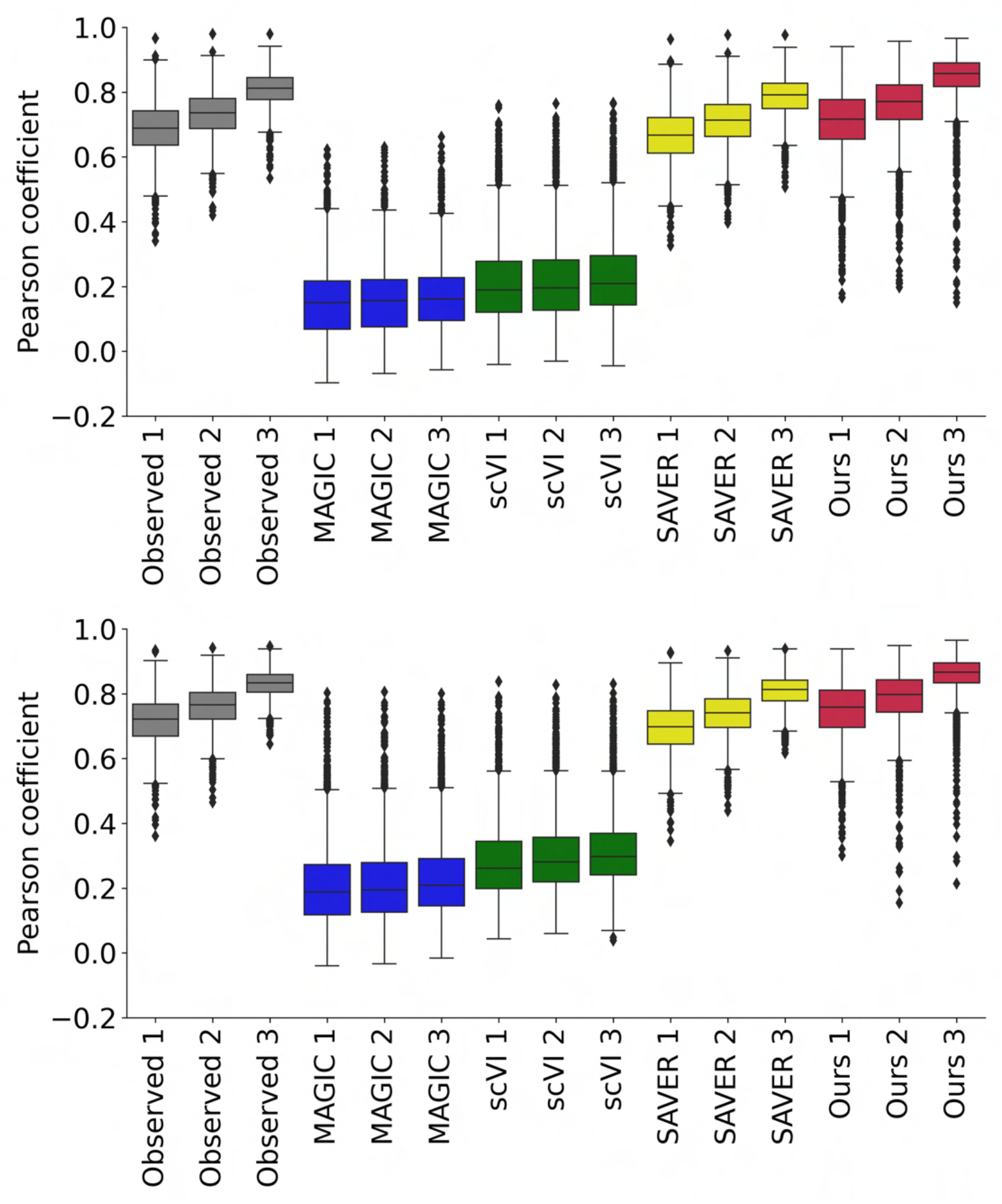
Box plots of Pearson coefficient between the reference and imputed data from different methods for EB differentiation data (first row) and zebrafish development data (second row). 1-3 shown in x-axis indicate the sampling efficiency 0.4%, 0.5%, and 0.75%.

## Discussion

Current scRNA-seq technologies suffer from a number of drawbacks, such as low capture efficiency, high dropout32, and measurement noise, and a preprocessing of the data is required for downstream applications. In this work, an effective TCER framework is proposed with incorporation of the gene-gene interactions of the system for accurate gene imputation. As revealed by the Pearson correlation analysis (Figs. 7 and 9), the existing techniques can hardly improve the data quality, which has recently been pointed out by Hou et al.^32^. TCER, on the other hand, greatly improves the Pearson correlation in all cases. Moreover, the proposed technique also improves the clustering and trajectory analysis substantially (Fig. 6, 5 and 8).

The TCER technique consists of two important steps. The GenoMap construction is critical in TCER for the network to recover the gene expression values. To illustrate this, we created a 2D map by randomly placing the genes on its grid and the map is processed by the same DNN in TCER for three different datasets. The Pearson correlations resulted from this random map + DNN is plotted in supplementary section 2. It is seen that the results are much inferior to that of GenoMap+DNN.

The construction of a GenoMap is less susceptible to noise and omitted gene expression values. This is attributed to the fact that the GenoMap construction depends on the distribution of expression values, instead of a particular gene expression value(s). Because of this, the TCER imputation results are robust even if some expression values are missing or noisy. The introduction of GenoMap helps the self-supervised DNN to learn the deep relationship among the genes and cells for better imputation.

The deep learning architecture in TCER adopts deformable convolutional operations for extraction of high-quality features^33^. As opposed to traditional convolution with a fixed kernel size, deformable convolution adaptively expands its receptive field by adjusting its kernel shape according to the input feature maps. Thus, the network can efficiently extract both low-level (local) and high-level (global) gene-gene interaction features. A skip connection between the downsampling and upsampling layers is introduced to preserve the multi-scale information. Finally, the dual-channel attention (channel-wise and pixel-wise attention) in our network can adaptively assign important feature information to important channels and positions, which allows the network to put emphasis on the important genes and differentially explore the gene-gene relationship for better data recovery.

In conclusion, we have proposed a novel TCER framework for gene expression recovery. The technique demonstrates unprecedented accuracy and reliability in gene imputation. Fundamentally different from the existing methods, TCER exploits the underlying gene-gene interaction information of a biological system via a transform-and-conquer strategy. TCER is self-supervised and its potential applications go much beyond genomic data processing. The same strategy should be applicable to other missing data problems in various biomedical and biocomputational applications.

## Methods

### Overview

As shown in Fig. 1, the proposed TCER consists of two steps: 1) GenoMap Construction and 2) GenoMap Recovery. Given a gene sequence data *S* ∈ ℝ^*q×m*^ where *q* is the number of cells while *m* is the number of genes in each cell, we first reconstruct the sequence data as 2D spatial data, termed as GenoMap, to represent the gene-gene relationship. Specifically, the generated GenoMap is denoted *G* ∈ ℝ^*q×s_w_×s_h_*^ where *s_w_, s_h_* is the width and height and q is the number of GenoMaps. Then, we design an end-to-end convolutional neural network named ER-Net for recovering gene expression. Due to the lack of ground-truth annotation, we introduce a novel self-supervised training strategy which simulates the dropout event of scRNA-seq data in the real world as self-supervision for learning the recovery of GenoMaps.

### GenoMap Construction

Intuition behind GenoMap Construction is to obtain a 2D spatial configuration of the genes where their interactive relationships could also be properly expressed. Further, genes with stronger interactions are supposed to have smaller Euclidean distances in the corresponding GenoMap. Specifically, in order to reposition the gene sequence data with m genes for each cell into a 2D GenoMap with a grid of *s_w_* × *s_h_* where *m* ≤ *s_w_* × *s_h_*, a pairwise interaction strength matrix *C* ∈ ℝ^*m×m*^ would first be calculated to maximize entropy of the whole sequence data. Then, a projection matrix *T* ∈ ℝ^*m*×^(*s_w_* × *s_h_*) is obtained to reconstruct the gene data into 2D grid based on maximally preserving the pairwise interactions. Specifically, pairwise interaction strength matrix could be calculated as follows^34^

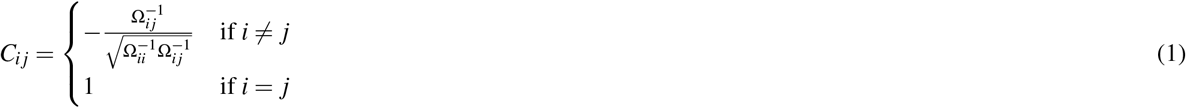

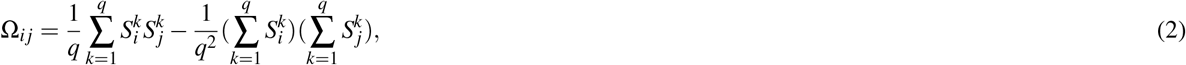

where Ω is the covariance matrix and 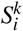 indicates the *i^th^* gene expression of the *k^th^* cell. Then Gromov-Wasserstein discrepancy between the pair interaction strength matrix *C* ∈ ℝ^*m×m*^ and the distance matrix 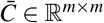 of the 2D grid space (*s_w_* × *s_h_*) is utilized based on its minimization to obtain the optimal projection matrix *T* ∈ ℝ^*m×*^(*s_w_* ×*s_h_*). The Gromov-Wasserstein discrepancy between matrices *C* and 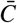 is defined as^35^

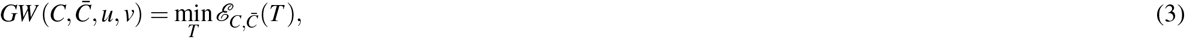

and

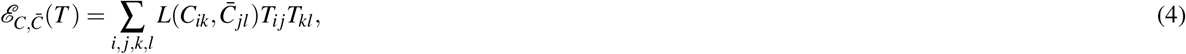

where *u, v* are vectors that contain relative importance of genes and locations in GenoMap. And *L* indicates the Kullback-Leibler divergence such that

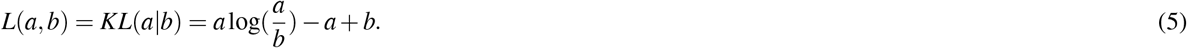

After the projection matrix is obtained, the restructured data could be written as

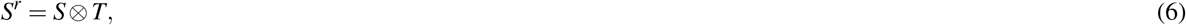

where *S^r^* ∈ ℝ^*q*x(*s_w_*×*s_h_*)^ and ⊗ represents the matrix multiplication. *S^r^* is then reshaped into the image format *G* ∈ ℝ^*q×S_w_×S_h_*^, i.e. the restructured GenoMap. And more detailed explanations could be found in ref.^36^.

### ER-Net

As shown in Fig.2, our proposed ER-Net follows a standard encoder-decoder structure^37^ where we plug in several cascaded DFA modules in the bottleneck to further learn global and local gene-gene interactions. In addition, we apply skip connections with channel-wise and spatial-wise attention to flexibly preserve information from shallow layers. Below we will give more details of ER-Net.

### Encoder-Decoder structure

In ER-Net, the encoder firstly produces feature maps at different resolutions by consecutively downsampling the 2D image features. Then three cascaded Deformable Fusion Attention (DFA) modules (see details in the Deformable Fusion Attention Module Section) are applied for exploring feature maps in low resolution with dynamic deformable receptive fields. Next, the decoder restores the 2D GenoMap at the original resolution by combining the upsampled features and the skip features from the encoder at different resolutions. Specifically, in the encoder, we use one convolutional layer with the stride of 1 and two convolutional layers with the stride of 2 for downsampling. And in the Decoder, we use two transposed convolutional layers with the stride of 2 and one convolutional layer with the stride of 1 for upsampling. The down/upsampling ratios are both 4×.

### Deformable Fusion Attention Module

Following the idea of fusion attention module^38^ and deformable convolution^33^, we design a Deformable Fusion Attention (DFA) module to better exploit the feature representation to recover 2D GenoMaps. As shown in Fig.2(b), each DFA module consists of two convolutional layers, one ReLU layer^39^, one deformable convolutional layer, Channel Attention and Pixel Attention. Different from the Fusion Attention (FA) module in FFA-Net^38^ which only includes the standard convolution with a fixed grid, here we employ the deformable convolutional layer^33^ to enable the deformation of the kernel shape. Such adaptive kernels can then expand the receptive field flexibly since the deformed grid sampling are combinations of various transformations e.g., scaling, rotation etc^33^. Specifically, for a standard convolution operation, the kernel shape is usually a fixed *n × n* square. Assume *n* is an odd integer. Denote the output feature map of the standard convolution as 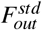. Then, in the *t^th^* layer, 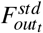 at position (*p_x_, p_y_*) could be described as

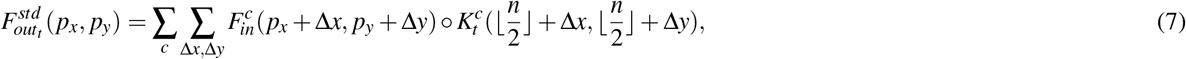

where ͦ denotes the Hadamard product, *c* enumerates all the channels of the input feature map *F_in_* and Δ*x*, Δ*y* indicates the offset of the sampled grid at position (*p_x_, p_y_*) where 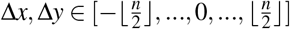. For the deformable convolution operation, the output feature map of the deformable convolution is denoted as 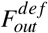. Then, in the *t^th^* layer, 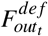 at position (*p_x_, p_y_*) can be described as

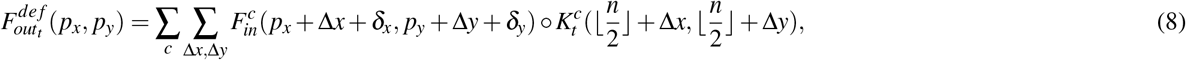

where *δ_x_, δ_y_* indicate the offsets to the fixed sampled grid in the standard convolution. These offsets are learned through standard convolution layers using *F_in_* as the input and optimized together with kernels during the training process. Therefore, by applying deformable convolution, features at any location inside this feature map could be considered altogether (*e.g*., (*p_x_, p_y_*)) instead of only the neighboring features, thus improving the representation capability of the network. Furthermore, we employ the channel attention module which aims to adaptively give different weights to distinct channels and the pixel attention module to adaptively give various weights to pixels at different locations. Such a dual-attention mechanism can automatically pay more attention to important features at different positions in the feature maps, which helps the network to capture more useful information. Residual learning is also included in the DFA module to prevent the degradation of the information^40^.

### Skip Connection with Attention

Inspired by Unet^41^, we have adopted skip connections between feature maps from upsampling and corresponding downsampling layers at different resolutions to enable the low-level information flow through the whole network which could then be beneficial for the GenoMap restoration. We also apply channel and spatial-wise attention mechanisms, following CBAM^42^ to adaptively assign higher weights to important channels and positions for fully exploiting features at different gene expression levels.

### Self-supervised training

Due to the lack of true gene expression data, in this paper, we propose a self-supervised training strategy by utilizing simulated dropout data as self-supervision for training the recovery network ER-Net. And we utilize *L*1 loss in both spatial and frequency domains during the training process for accurate restorations.

### Dropout Simulation for Self-supervision

Inspired by the fact that real-world scRNA-seq data usually suffers from high dropouts^43^, we propose a self-supervised strategy which utilizes *Dropout Simulation* to imitate the dropout events during training. Specifically, it essentially applies random masking to generate the input data for self-supervision. Such a self-supervised strategy not only effectively reduces the need for the ground-truth annotation but also helps ER-Net to learn the common representation among the GenoMaps. Specifically, given a batch of GenoMap 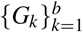 with *b* as the batch size, for each GenoMap, a dropout ratio *r_k_* is randomly selected among [*r^l^, r^u^*] at each training step. And *r^l^, r^u^* ∈ [0,1] are both fractions indicating the lower and upper bound respectively. Suppose at training step *s*, the randomly selected dropout ratio for *G_k_* is denoted as 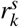. The pixel indices of non-zero values inside *G_k_* are denoted as 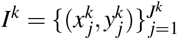 where *J^k^* is the number of non-zero values in *G_k_* and 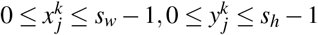. Then, a subset 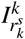 is randomly selected from *I^k^* based on 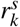 where the number of pixel indexes in 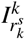 is 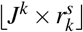. And a placeholder *h* is then used to replace the selected non-zero values indicating the masked positions that

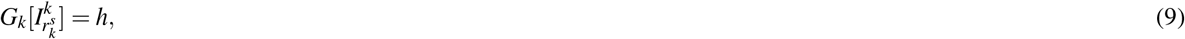

where *h* ∈ [0,1] is a fraction number. Besides, since scRNA-seq data is always sparse and has a lot true zeros, to help the network learn to distinguish those true zeros from dropout events, we also use *h* to replace all zero values in *G_k_* that

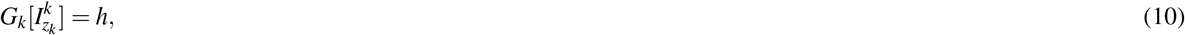

where 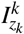 indicates all the pixel indexes with zero values in *G_k_*. And the corresponding dropout GenoMap at training step *s* is denoted as 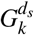 and the corresponding random dropout simulation process is then described as

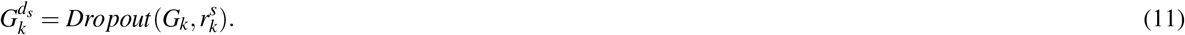

By applying this random dropout strategy, ER-Net could learn some shared patterns among all the GenoMap which then contribute to better data recovery.

### Overall training loss

We take both spatial and frequency domain information into consideration for the overall objective function. *L*1 loss in the spatial domain is applied to ensure the accurate recovery for each pixel in GenoMap while we employ the frequency loss to better restore the high frequency information as well. Since the goal of the proposed ER-Net is to recover *G_k_* based on the input 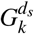, thus, according to the overall loss, the optimization objective could be described as

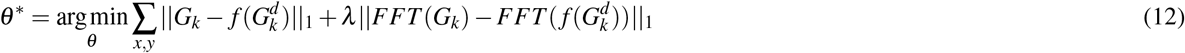

where *f* indicates the ER-Net with *θ* as its parameters, *x* ∈ [0,*s_w_* −1],*y* ∈ [0,*s_h_* −1], *l* is the weight for the frequency loss and *FFT* indicates fast Fourier transform function. Thus, for overall loss at training step *s*, we have

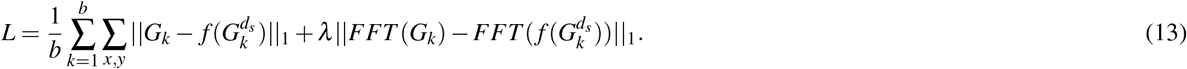

### Generation of observation datasets

Following SAVER^12^, we first select high-quality genes and cells based on their expression levels and refer to them as the true expression *λ_gc_*. The observed gene sequence dataset *S* is then constructed by applying Poisson–gamma mixture on *λ_gc_* which is also known as the negative binomial model. Specifically, *τ_c_* is sampled from a gamma distribution simulating the uncertainty of the gene values and a Poisson distribution is then placed on *τ_c_λ_gc_* to get the simulated observation *S* which could be described as

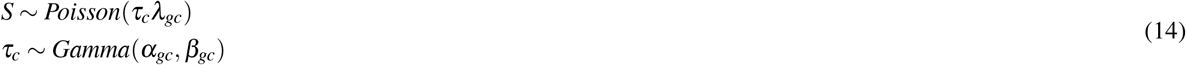

where *α_gc_, β_gc_* are the shape and rate parameters respectively that control the mean and variance. And *τ_c_* could be considered as the efficiency loss from the true expression data *λ_gc_*.

### Implementation Details

GenoMap construction is implemented in MATLAB while the imputation experiments are implemented on one RTX2080 GPU in PyTorch. For the imputation network, we adopt Adam optimizer for network parameters optimization, setting the learning rate as 0.0002, *β*_1_ as 0.9, *β*_2_ as 0.999 and *ε* as 10^-8^. Batch size is set as 64 and the learning rate is half decayed every 40 epochs. And we train our ER-Net for 100 epochs. And the placeholder *h* is set as 0.999.

Moreover, to mimic variation in efficiency across cells, *τ_c_* is sampled as follows^12^

- 0.75% efficiency *τ*_*c*1_ ~ Gamma(15, 2000)
- 0.5% efficiency *τ*_*c*2_ ~ Gamma(10, 2000)
- 0.4% efficiency *τ*_*c*3_ ~ Gamma(8, 2000)

### Datasets

We evaluate our proposed method on six different real-world datasets. For each dataset, we generate three different observation datasets with different efficiency loss. And Pearson coefficient is adopted to evaluate the performance. Pearson coefficient is calculated for each gene across the whole cells and is analyzed on the reference data (true expression), the observed data (input), and the imputed data (output). Details of the six real-world datasets are discussed below.

### Mouse intestinal epithelium

Intestinal epithelial cells absorb nutrients, respond to microbes, function as a barrier and help to coordinate immune responses. Original dataset includes 53,193 individual epithelial cells from the small intestine and organoids of miceet al.^25^, which enabled the identification and characterization of previously unknown intestinal epithelial cell subtypes and their genetic characteristics. In our experiments, we select 4072 cells and 1776 genes in total as our dataset.

### Engineered 3D neural tissues

Human engineered neural tissues are very helpful in understanding neurological diseases. Human neural cells from this dataset^26^ are cultured with differentiated human astrocytic cells where human embryonic stem cells (hESC) induced neuronal cells and human astrocytic cells differentiated from hESCs were co-cultured at 1:1 ratio in a 3D composite hydrogel. In our study, we choose a subset of 2364 cells and 2735 genes from the original dataset.

### Mammalian brains

A highly scalable single-nucleus RNA-seq (sNucDrop-seq) approach^27^ is developed for massively parallel scRNA-seq without enzymatic dissociation and nucleus sorting. This dataset is acquired by sNucDrop-seq. Such technology could accurately resolve cellular diversity in a low-cost and high-throughput manner and provide an unbiased isolation of intact single cells from complex tissues such as adult mammalian brains. In the original dataset^27^, 18,194 nuclei were isolated from cortical tissues of adult mice. With extensive evaluations, the authors illustrate that sNucDrop-seq not only reveals neuronal and non-neuronal subtype composition with high accuracy but can also analyze transient transcriptional states driven by neuronal activity at single cell resolution. We select 10360 cells and 2344 genes from the original dataset in our experiments.

### Cellular taxonomy of the mouse bone marrow stroma

Stroma plays an important role in the development, homeostasisment and repair of organs. This dataset defines cellular taxonomy of the mouse bone marrow stroma and its perturbation by malignancy using scRNA-seq^28^. Seventeen stromal subsets were identified expressing distinct hematopoietic regulatory genes spanning new fibroblastic and osteoblastic subpopulations including distinct osteoblast differentiation trajectories. Emerging acute myeloid leukemia impaired mesenchymal osteogenic differentiation and reduced regulatory molecules necessary for normal hematopoiesis. This taxonomy of the stromal compartment provides a comprehensive bone marrow cell census and experimental support for cancer cell crosstalk with specific stromal elements that impair normal tissue function and thus lead to new cancers. In our study, we use a subset of 12162 cells with 2422 genes for each cell as our dataset.

### Embryoid bodies

EB differentiation outlines the key aspects of early embryogenesis and has been successfully used as the first step in differentiation protocols for certain types of neurons, astrocytes and oligodendrocytes, hematopoietic, endothelial and muscle cells, hepatocytes and pancreatic cells, and germ cells. Approximately 31,000 cells were measured in this dataset where they were evenly distributed over a 27-day differentiation time course^29^. And samples were collected at 3-d intervals and pooled for measurement on the 10x Chromium platform^29,31^. In our experiments, we selected 9754 cells with 2282 genes for each cell as our dataset.

### Zebrafish embryogenesis

To reveal the transcriptional trajectories during the development of zebrafish embryos, this dataset^30^ was profiled using the scRNA-seq technology called Drop-seq. It includes 38,731 cells from 694 embryos across 12 closely spaced stages of early zebrafish development. Data were acquired from the high blastula stage (3.3 hours postfertilization, moment after transcription starts from the zygotic genome) to six-somite stage (12 hours after postfertilization, just after gastrulation). Due to the pluripotency at high blastula stage, most of the cells then have differentiated into specific cell types at six-somite stage. And we use 20014 cells and 2341 genes as our dataset in our study.

## Data and code availability

The datasets generated during the current study, TCER imputation results along with the corresponding checkpoint and the implementation codes are all publicly available at https://anonymous.4open.science/r/TCER-37E7.

## References

1. Qiu, X. et al. Single-cell mrna quantification and differential analysis with census. Nat. methods 14, 309–315 (2017).

2. Vu, T. N. et al. Beta-poisson model for single-cell rna-seq data analyses. Bioinforma. 32, 2128–2135 (2016).

3. Miao, Z., Deng, K., Wang, X. & Zhang, X. Desingle for detecting three types of differential expression in single-cell rna-seq data. Bioinforma. 34, 3223–3224 (2018).

4. Kiselev, V. Y. et al. Sc3: consensus clustering of single-cell rna-seq data. Nat. methods 14, 483–486 (2017).

5. Xu, C. & Su, Z. Identification of cell types from single-cell transcriptomes using a novel clustering method. Bioinforma.31, 1974–1980 (2015).

6. Lin, P., Troup, M. & Ho, J. W. Cidr: Ultrafast and accurate clustering through imputation for single-cell rna-seq data. Genome biology 18, 1–11 (2017).

7. Trapnell, C. et al. The dynamics and regulators of cell fate decisions are revealed by pseudotemporal ordering of single cells. Nat. biotechnology 32, 381–386 (2014).

8. Setty, M. et al. Wishbone identifies bifurcating developmental trajectories from single-cell data. Nat. biotechnology 34, 637–645 (2016).

9. Street, K. et al. Slingshot: cell lineage and pseudotime inference for single-cell transcriptomics. BMC genomics 19, 1–16 (2018).

10. Welch, J. D., Hu, Y. & Prins, J. F. Robust detection of alternative splicing in a population of single cells. Nucleic acids research 44, e73–e73 (2016).

11. Huang, Y. & Sanguinetti, G. Brie: transcriptome-wide splicing quantification in single cells. Genome biology 18, 1–11 (2017).

12. Huang, M. et al. Saver: gene expression recovery for single-cell rna sequencing. Nat. methods 15, 539–542 (2018).

13. Wang, J. et al. Data denoising with transfer learning in single-cell transcriptomics. Nat. methods 16, 875–878 (2019).

14. Tang, W. et al. baynorm: Bayesian gene expression recovery, imputation and normalization for single-cell rna-sequencing data. Bioinforma. 36, 1174–1181 (2020).

15. Li, W. V. & Li, J. J. An accurate and robust imputation method scimpute for single-cell rna-seq data. Nat. communications9, 1–9 (2018).

16. Van Dijk, D. et al. Recovering gene interactions from single-cell data using data diffusion. Cell 174, 716–729 (2018).

17. Gong, W., Kwak, I.-Y., Pota, P., Koyano-Nakagawa, N. & Garry, D. J. Drimpute: imputing dropout events in single cell rna sequencing data. BMC bioinformatics 19, 1–10 (2018).

18. Talwar, D., Mongia, A., Sengupta, D. & Majumdar, A. Autoimpute: Autoencoder based imputation of single-cell rna-seq data. Sci. reports 8, 1–11 (2018).

19. Arisdakessian, C., Poirion, O., Yunits, B., Zhu, X. & Garmire, L. X. Deepimpute: an accurate, fast, and scalable deep neural network method to impute single-cell rna-seq data. Genome biology 20, 1–14 (2019).

20. Lopez, R., Regier, J., Cole, M. B., Jordan, M. I. & Yosef, N. Deep generative modeling for single-cell transcriptomics. Nat. methods 15, 1053–1058 (2018).

21. Deng, Y., Bao, F., Dai, Q., Wu, L. F. & Altschuler, S. J. Scalable analysis of cell-type composition from single-cell transcriptomics using deep recurrent learning. Nat. methods 16, 311–314 (2019).

22. Zappia, L., Phipson, B. & Oshlack, A. Splatter: simulation of single-cell rna sequencing data. Genome biology 18, 1–15 (2017).

23. Van der Maaten, L. & Hinton, G. Visualizing data using t-sne. J. machine learning research 9 (2008).

24. McInnes, L., Healy, J. & Melville, J. Umap: Uniform manifold approximation and projection for dimension reduction. arXiv preprint arXiv:1802.03426 (2018).

25. Haber, A. L. et al. A single-cell survey of the small intestinal epithelium. Nat. 551, 333–339 (2017).

26. Tekin, H. et al. Effects of 3d culturing conditions on the transcriptomic profile of stem-cell-derived neurons. Nat. biomedical engineering 2, 540–554 (2018).

27. Hu, P. et al. Dissecting cell-type composition and activity-dependent transcriptional state in mammalian brains by massively parallel single-nucleus rna-seq. Mol. cell 68, 1006–1015 (2017).

28. Baryawno, N. et al. A cellular taxonomy of the bone marrow stroma in homeostasis and leukemia. Cell 177, 1915–1932 (2019).

29. Martin, G. R. & Evans, M. J. Differentiation of clonal lines of teratocarcinoma cells: formation of embryoid bodies in vitro. Proc. Natl. Acad. Sci. 72, 1441–1445 (1975).

30. Farrell, J. A. et al. Single-cell reconstruction of developmental trajectories during zebrafish embryogenesis. Sci. 360, eaar3131 (2018).

31. Moon, K. R. et al. Visualizing structure and transitions in high-dimensional biological data. Nat. biotechnology 37, 1482–1492 (2019).

32. Hou, W., Ji, Z., Ji, H. & Hicks, S. C. A systematic evaluation of single-cell rna-sequencing imputation methods. Genome biology 21, 1–30 (2020).

33. Dai, J. et al. Deformable convolutional networks. In Proceedings of the IEEE international conference on computer vision, 764–773 (2017).

34. Stein, R. R., Marks, D. S. & Sander, C. Inferring pairwise interactions from biological data using maximum-entropy probability models. PLoS computational biology 11, e1004182 (2015).

35. Peyré, G., Cuturi, M. & Solomon, J. Gromov-wasserstein averaging of kernel and distance matrices. In International Conference on Machine Learning, 2664–2672 (PMLR, 2016).

36. Islam, M. T. & Xing, L. Cartography of genomic interactions enables deep analysis of single-cell expression data. Nat. communications 14, 679 (2023).

37. Cho, K. et al. Learning phrase representations using rnn encoder-decoder for statistical machine translation. arXiv preprint arXiv:1406.1078 (2014).

38. Qin, X., Wang, Z., Bai, Y., Xie, X. & Jia, H. Ffa-net: Feature fusion attention network for single image dehazing. In Proceedings of the AAAI Conference on Artificial Intelligence, vol. 34, 11908–11915 (2020).

39. Agarap, A. F. Deep learning using rectified linear units (relu). arXiv preprint arXiv:1803.08375 (2018).

40. He, K., Zhang, X., Ren, S. & Sun, J. Deep residual learning for image recognition. In Proceedings of the IEEE conference on computer vision and pattern recognition, 770–778 (2016).

41. Ronneberger, O., Fischer, P. & Brox, T. U-net: Convolutional networks for biomedical image segmentation. In International Conference on Medical image computing and computer-assisted intervention, 234–241 (Springer, 2015).

42. Woo, S., Park, J., Lee, J.-Y. & Kweon, I. S. Cbam: Convolutional block attention module. In Proceedings of the European conference on computer vision (ECCV), 3–19 (2018).

43. Haque, A., Engel, J., Teichmann, S. A. & Lönnberg, T. A practical guide to single-cell rna-sequencing for biomedical research and clinical applications. Genome medicine 9, 1–12 (2017).

