## Supplemental Table 1 for "Self-supervised deep learning of gene-gene interactions for improved gene expression recovery"

### GenoMap provides better space configurations for gene expression recovery

GenoMap offers a better way to reposition the gene expression data and the space configurations of GenoMap explicitly takes the gene-gene relationship into consideration which contributes to the imputation results. To demonstrate this, we adopt an ablation study which reshapes the raw gene data into 2D format without taking their interactions into consideration as the input of our proposed ER-Net. To be specific, given a gene sequence data $S$ with $q$ cells and $m$ genes, unlike constructing $S$ into GenoMap with the optimized projection matrix, each cell is directly reshaped into an image format $(s_w \times s_h)$ where $m = s_w \times s_h$. And these reshaped images would then become the inputs of ER-Net. We particularly include one real-world dataset (cellular taxonomy data) and apply three different efficiency loss to get the corresponding observation datasets. We use **w/ GenoMap** to indicate using GenoMap as the inputs for imputation while **w/o GenoMap** to represent imputing the directly reshaped images. As shown in Table 1, we could notice that for all observations, **w/ GenoMap** all outperform **w/o GenoMap** (0.7855 vs. 0.7693, 0.8218 vs. 0.8094 and 0.8608 vs. 0.8431). This indicates that after repositioning the data, GenoMap has a better configured space compared with directly reshaped images and thus, could contribute to the better imputation results.

| | $\tau_{c1}$ | $\tau_{c2}$ | $\tau_{c3}$ |
|---|---|---|---|
| w/o GenoMap | 0.7693 ± 0.1125 | 0.8094 ± 0.1090 | 0.8431 ± 0.1067 |
| w/ GenoMap | 0.7855 ± 0.1077 | 0.8218 ± 0.0973 | 0.8608 ± 0.0868 |

**Table 1.** Ablation study on the effect of GenoMap construction in our gene data recovery pipeline. Experiments are conducted on three observation datasets which are sampled at different efficiency loss from cellular taxonomy data.
